# Repurposing Quetiapine as an Adjuvant Therapeutic Agent for Triple-Negative Breast Cancer

**DOI:** 10.1101/2025.04.24.650517

**Authors:** Ling He, Purva Joshi, Angeliki Ioannidis, Anoushka Kathiravan, Mohammad Saki, Frank Pajonk

**Author notes:** **Correspondence address:** Ling He, PhD, Department of Radiation Oncology, David Geffen School of Medicine at UCLA, 10833 Le Conte Ave, Los Angeles, CA 90095-1714.

## Abstract

**Purpose:** Triple-negative breast cancer (TNBC) lacks actionable molecular targets, so treatment primarily relies on cytotoxic chemotherapy and radiotherapy, yet relapse, resistance, and metastasis still drives poor long-term survival. Dopamine signaling has recently been linked to tumor aggressiveness, and dopaminel1lreceptor antagonists have shown preclinical benefit in other cancers. We therefore evaluated quetiapine (QTP), an FDAl1lapproved DRD2/DRD3 antagonist, as a therapeutic adjunct in TNBC.

**Methods:** SUM159l1lPT, BTl1l549, and MDAl1lMBl1l231 cells were treated with QTP alone or combined with radiation or standard agents (doxorubicin, paclitaxel, 5l1lfluorouracil). Clonogenic and mammosphere assays measured proliferative and selfl1lrenewal potential. Annexin V/propidiuml1liodide flow cytometry quantified apoptosis, and γl1lH2AX immunofluorescence tracked DNA doublel1lstrand breaks and repair kinetics. Transwell assays assessed migration of untreated bulk cells and radiationl1lsurviving subclones.

**Results:** QTP significantly reduced clonogenicity and self-renewal in all TNBC models tested, both alone and in combination with radiation or chemotherapy. In apoptosis assays, QTP treatment induced a marked increase in early and late apoptotic cell populations. QTP also promoted DNA double-strand break formation and delayed repair, as indicated by persistent γ-H2AX foci at 24 hours post-treatment. Additionally, QTP impaired the migratory capacity of both untreated and radiation-surviving cells. Combination treatments with QTP and doxorubicin produced synergistic effects, resulting in complete loss of colony-forming ability and mammosphere formation.

**Conclusion:** The data presented support the repurposing of quetiapine as an adjuvant therapeutic agent alongside radiotherapy and/or chemotherapy in TNBC. By targeting apoptosis, DNA repair, and cancer cell migration, QTP offers a novel, multi-faceted approach to improve outcomes in this high-risk breast cancer subtype.

## Introduction

Breast cancer remains the most commonly diagnosed cancer among women, accounting for approximately 30 % (or 1 in 3) of all new female cancer cases each year. Among these, triple-negative breast cancer (TNBC) constitutes an estimated 10 - 15 % of all cases, which translates to roughly 30,000 to 45,000 new diagnoses annually in the United States [1, 2] (https://www.nationalbreastcancer.org/breast-cancer-facts/). TNBC is defined by a lack of estrogen receptor (ER), progesterone receptor (PR), and human epidermal growth factor receptor 2 (HER2), that are commonly used as therapeutic targets in other breast cancer subtypes. Due to this absence of targetable biomarkers, TNBC has been recognized as one of the most aggressive and therapeutically challenging forms of breast cancer [3–5].

Despite continued advances in early detection and prevention strategies, the American Cancer Society predicts that the number of newly diagnosed invasive breast cancer cases is expected to rise further in 2025 (https://www.cancer.org/cancer/types/breast-cancer/about/how-common-is-breast-cancer.html), highlighting the persistent burden of this disease. TNBC is associated with rapid tumor progression, increased rates of distant metastasis, early relapse -often within the first three years of diagnosis- and significantly worse overall survival compared to other forms of breast cancer [6, 7]. Current standard-of-care relies heavily on cytotoxic chemotherapy and radiotherapy, as targeted hormonal and HER2-directed therapies are ineffective in this case [8, 9]. However, the clinical benefit of chemotherapy and/or radiotherapy is often limited by TNBC’s pronounced intra-tumoral heterogeneity, high proliferative rate, and frequent enrichment of therapy-resistant cancer stem-like cells, which attribute to treatment failure and recurrence [10, 11]. Collectively, these challenges underscore an urgent and unmet need for the development of novel therapeutic strategies that can effectively overcome treatment resistance, eradicate tumor-initiating cells, and ultimately improve long-term outcomes for patients with this aggressive disease subtype.

Dopamine is a well-characterized neurotransmitter that plays a central role in regulating numerous complex brain functions, including mood, motivation, cognition, and motor control [12, 13]. Interestingly, beyond its neurological roles, emerging evidence has highlighted dopamine signaling as a potential modulator in cancer biology [14–18]. In particular, the D2-like class of dopamine receptors -which includes DRD2, DRD3, and DRD4-has been found to be upregulated in various malignancies and linked to cancer cell stemness [19–21]. Moreover, reduced cancer risk has been correlated with disorders such as schizophrenia and Parkinson’s disease [19, 22], in which dopaminergic drugs are used, suggesting a potential protective role of dopamine receptor antagonism in cancer development.

Quetiapine (QTP), an FDA-approved atypical antipsychotic, functions primarily as an antagonist of dopamine receptors D2 and D3 (DRD2/DRD3) and is widely used in the clinical management of schizophrenia and bipolar disorder. In our previous work, we demonstrated the therapeutic potential of QTP in treating glioblastoma, one of the most aggressive brain tumors, by targeting therapy-resistant cancer stem-like cells [21]. Based on these findings, we sought to investigate whether QTP could exert similar anti-tumor effects when used in combination with radiotherapy and/or chemotherapeutic agents in TNBC.

## Materials and Methods

### TCGA data mining and analysis

Data from The Cancer Genome Atlas (TCGA) Invasive Breast Carcinoma cohort (assessed via cBioPortal on February 10, 2025) was used for this analysis. Expression data (RNA-Seq V2 RSEM Z scores) and overall survival data for DRD2 and DRD3 were retrieved. Only patients with available data for both gene expression and survival were included in the analysis (N=1980). Kaplan-Meier survival analysis was performed using a Z-score cut-off of 1.0, and the patients were stratified into subgroups based on expression levels: up-regulated, down-regulated, and unchanged. These stratified groups were then used to estimate overall survival differences.

To specifically investigate the prognostic relevance of DRD2 in TNBC, we further filtered the dataset to include only patients with confirmed ER-negative, PR-negative, and HER2-negative status (N=288). Expression data and survival information for DRD2 were extracted using the Z-score cut-off of 1.5. Kaplan-Meier survival analysis was then conducted to evaluate the association between DRD2 expression and overall survival specifically within the TNBC subset.

### Cell lines

Human triple-negative breast cancer cell line SUM159-PT (RRID:CVCL_5423) was purchased from Asterand (Detroit, MI) and cultured in F12 Medium (Invitrogen, Carlsbad, CA) supplemented with 5% fetal bovine serum (FBS), penicillin (100 units/ml), streptomycin (100 μg/ml), 5 μg/ml insulin (Eli Lilly, Indianapolis, IN), 0.1% 1M HEPES buffer (Invitrogen) and 1 μg/ml hydrocortisone (Pfizer, New York, NY), under log-phase growth conditions. Human triple-negative breast cancer cell lines BT-549 (RRID:CVCL_1092) and MDA-MB-231 (RRID:CVCL_0062) were obtained from American Type Culture Collection (ATCC). MDA-MB-231 cells were cultured in Dulbecco’s Modified Eagle Medium (DMEM; Thermo Fisher) with 10% FBS, and penicillin-streptomycin, while BT-549 cells were maintained in RPMI-1640 medium (Thermo Fisher) supplemented with 10% FBS, 1% penicillin-streptomycin, and insulin (0.023 IU/ml). All cell lines were grown in a humidified incubator at 37°C with 5% CO2 and routinely tested for mycoplasma infection (#G238, Applied biological Materials, Ferndale, WA). The unique identity of all cell line was confirmed by DNA fingerprinting (Laragen, Culver City, CA).

### Drug treatment

Quetiapine (#KS-1099, Key Organic) was dissolved at a stock concentration of 10 mM in DMSO, and 1 µM, 2.5 µM, 5 µM, 10 µM, 20 µM were used. Paclitaxel (#S1150, Selleckchem) was dissolved at a stock concentration of 5 mM in DMSO, and 1 nM, 5 nM, 10 nM were used. 5-fluorouracil (#S1209, Selleckchem) was dissolved at a stock concentration of 10 mM in DMSO, and 0.5 µM, 1 µM, 5 µM were used. Doxorubicin (#15007, Cayman Chemical) was dissolved at a stock concentration of 5 mM in DMSO, and 5 nM, 10 nM, 20 nM were used.

### Quantitative Reverse Transcription-PCR

Total RNA was isolated using TRIZOL Reagent (Invitrogen). cDNA synthesis was carried out using the SuperScript Reverse Transcription IV (Invitrogen). Quantitative PCR was performed in the QuantStudio^TM^ 3 Real-Time PCR System (Applied Biosystems, Carlsbad, CA, USA) using the PowerUp^TM^ SYBRTM Green Master Mix (Applied Biosystems). C_t_ for each gene was determined after normalization to GAPDH and ΔΔC_t_ was calculated relative to the designated reference sample. Gene expression values were then set equal to 2^−ΔΔCt^ as described by the manufacturer of the kit (Applied Biosystems). All PCR primers were synthesized by Invitrogen with PPIA as the housekeeping gene (for primer sequences see **Supplemental Table 1**).

### Clonogenic survival assay

SUM159-PT, BT-549, and MDA-MB-231 cells were trypsinized and plated into 6-well plates at a density of 100-200 cells per well. The following day, cells were treated with either DMSO, QTP, chemotherapeutic drugs or QTP plus chemotherapeutic drugs at the indicated concentrations. Culture media was refreshed every three days. After two weeks, colonies were fixed and stained with 0.5% crystal violet. Colonies consisting of at least 50 cells were counted in each group and presented as percentage of colonies formed relative to the number of cells initially plated.

### Mammosphere Formation Assay with Extreme Limiting Dilution Analysis (ELDA)

SUM159-PT and BT-549 mammospheres were dissociated by TrypLE™ and plated into 96-well ultra-low attachment plates in mammosphere media (DMEM-F12, 10 ml/500 ml B27 [Invitrogen], 5 µg/ml bovine insulin [Sigma], 4 µg/ml heparin [Sigma], 20 ng/ml basic fibroblast growth factor [bFGF, Sigma], and 20 ng/ml epidermal growth factor [EGF, Sigma]). For ELDA, cells were seeded at varying densities: 1 to 512 cells per well for 0 Gy conditions, 2 to 1024 cells per well for 4 Gy, and 4 to 2048 cells per well for 8 Gy. Cells were treated with either QTP (10 µM) or DMSO solvent control one hour prior to receiving a single dose of 0, 4, or 8 Gy radiation the next day. Growth factors (EGF and bFGF) were replenished every 3 days. Mammospheres were cultured for 2-3 weeks, after which wells containing spheres were counted. Results were expressed as the percentage of wells with sphere formation relative to the initial number of cells plated. The stem cell frequency was calculated using the ELDA software (http://bioinf.wehi.edu.au/software/elda/) [23].

### Irradiation

Cells were irradiated at room temperature using an experimental X-ray irradiator (Gulmay Medical Inc. Atlanta, GA) at a dose rate of 5.519 Gy/min. The X-ray beam was operated at 300 kV and hardened using a 4 mm Be, a 3 mm Al, and a 1.5 mm Cu filter and calibrated using NIST-traceable dosimetry. Corresponding controls were sham irradiated.

### Annexin V/PI Staining and Flow Cytometry

SUM159-PT and BT-549 monolayer cells were plated onto 10-cm petri dishes and treated with QTP (10 µM) one hour prior to receiving a single dose of 4 Gy irradiation the following day. 72 hours post-irradiation, cells were harvested, washed twice with cold PBS, and resuspended in 1X Annexin V binding buffer at a concentration of 1 × 10^6^ cells/ml. For each sample, 100 μl of cell suspension was incubated with 5 μl of FITC-conjugated Annexin V and 5 μl of propidium iodide (PI, 50 μg/ml) for 15 minutes at room temperature in the dark. Following incubation, 400 μl of binding buffer was added, and samples were immediately analyzed using a BD LSR Fortessa flow cytometer (BD Bioscience, San Jose, CA), acquiring a minimum of 50,000 events per sample. Data analysis were performed using FlowJo v10 software. Cells positive for Annexin V and negative for PI were defined as early apoptotic, double-positive cells as late apoptotic or necrotic, and double-negative cells are viable cells.

### Cell Proliferation

SUM159-PT and BT-549 monolayers were trypsinized and seeded at a density of 10,000 cells per 4-cm petri dish. The following day, cells were treated with either DMSO or QTP (10 µM), with or without a single dose of 4 Gy radiation. Cell proliferation was assessed by harvesting and counting cells on day 5, 10, 15, and 20 post-treatment.

### Migration/Invasion Assay

Serum-starved SUM159-PT or BT-549 monolayers were trypsinized and plated onto Transwell inserts (#08-771-12, Fisher Scientific) at a density of 1 x 10^5^ cells per well. QTP or solvent control (DMSO) was added to the medium in the bottom chamber containing 10% FBS. A negative control using 1% FBS medium was included to eliminate the FBS gradient between the upper and lower chambers. After 16-24 hours of incubation, cells were fixed with 10% formalin. Non-migrated cells on the upper side of the membrane were gently removed with cotton swabs, while migrated cells on the lower side were stained with 0.5% crystal violet. Images were captured using a digital microscope (BZ-9000, Keyence, Itasca, IL) and quantified using Image J software.

For experiments involving irradiation, SUM159-PT or BT-549 monolayers were plated onto 10-cm Petri dishes and treated with 10 µM QTP, with or without a single dose of 4 Gy irradiation. 48 hours post-irradiation, the cells were serum starved overnight. The following day, cells were trypsinized and plated onto Transwell inserts at 1 x 10^5^ cells per well. The bottom chamber contained 10% FBS medium to create an FBS gradient that promotes cell migration. After 16-24 hours, cells were fixed and stained with crystal violet as described above.

#### γ-H2AX immunofluorescence staining

SUM159-PT monolayer cells were seeded onto the round glass coverslips at a density of 1 x 10^5^ cells/well in the 6-well plate. The following day, cells were treated with DMSO or QTP (10 µM), with or without a single dose of 4 Gy radiation. 6 hours and 24 hours post-treatment, the coverslips were fixed, permeabilized, and blocked with 10% goat serum in PBS for 1 hour at RT. The γ-H2AX primary antibody (Cell Signaling Technology, #9718s, 1:250) was then applied and incubated overnight at 4°C. The next day, the secondary antibodies Alexa Fluor 594 Goat Anti-rabbit immunoglobulin G (IgG) (H/L) antibody (1:1,000 (Invitrogen)) was applied for 60 min, with subsequent nuclear counterstaining with Hoechst 33342 (Invitrogen, Cat# H3570, 1:5000). The sections were sealed with VECTASHIELD^®^ PLUS Antifade Mounting Medium (Vector Laboratories, Cat# H-1900) and images were taken with a digital microscope (BZ-9000, Keyence, Itasca, IL).

### Statistics

All data are represented as mean ± standard error mean of at least 3 biologically independent experiments. A *P* value of < 0.05 in a 2-sided student t-test or ANOVA indicated a statistically significant difference.

## Results

### Up-regulated DRD2 expression is associated with decreased overall survival in invasive breast cancer patients

To determine whether the expression levels of dopamine receptors DRD2 and DRD3 correlate with clinical outcomes in invasive breast cancer, we analyzed overall survival (OS) data from 1,980 invasive breast cancer patients within the TCGA Provisional dataset. Patients were stratified into three subgroups based on DRD2 and DRD3 expression levels (upregulated, downregulated, or unchanged), using a Z-score cutoff of ±1.0. Our analysis revealed that patients exhibiting upregulated DRD2 expression demonstrated significantly poorer median overall survival (111.10 months; *p* = 0.0245) compared to patients with unchanged DRD2 expression (157.83 months) **(Figure 1A/B)**. In contrast, the median survival for patients with downregulated DRD2 expression (151.17 months) did not significantly differ from patients with normal expression (*p* = 0.801) **(Supplemental Figure 1A/B)**. Furthermore, no significant difference in median survival was observed between patients with either upregulated or downregulated DRD3 expression and those with unchanged DRD3 expression levels **(Figure 1C/D; Supplemental Figure 1C/D)**.

**Figure 1.**
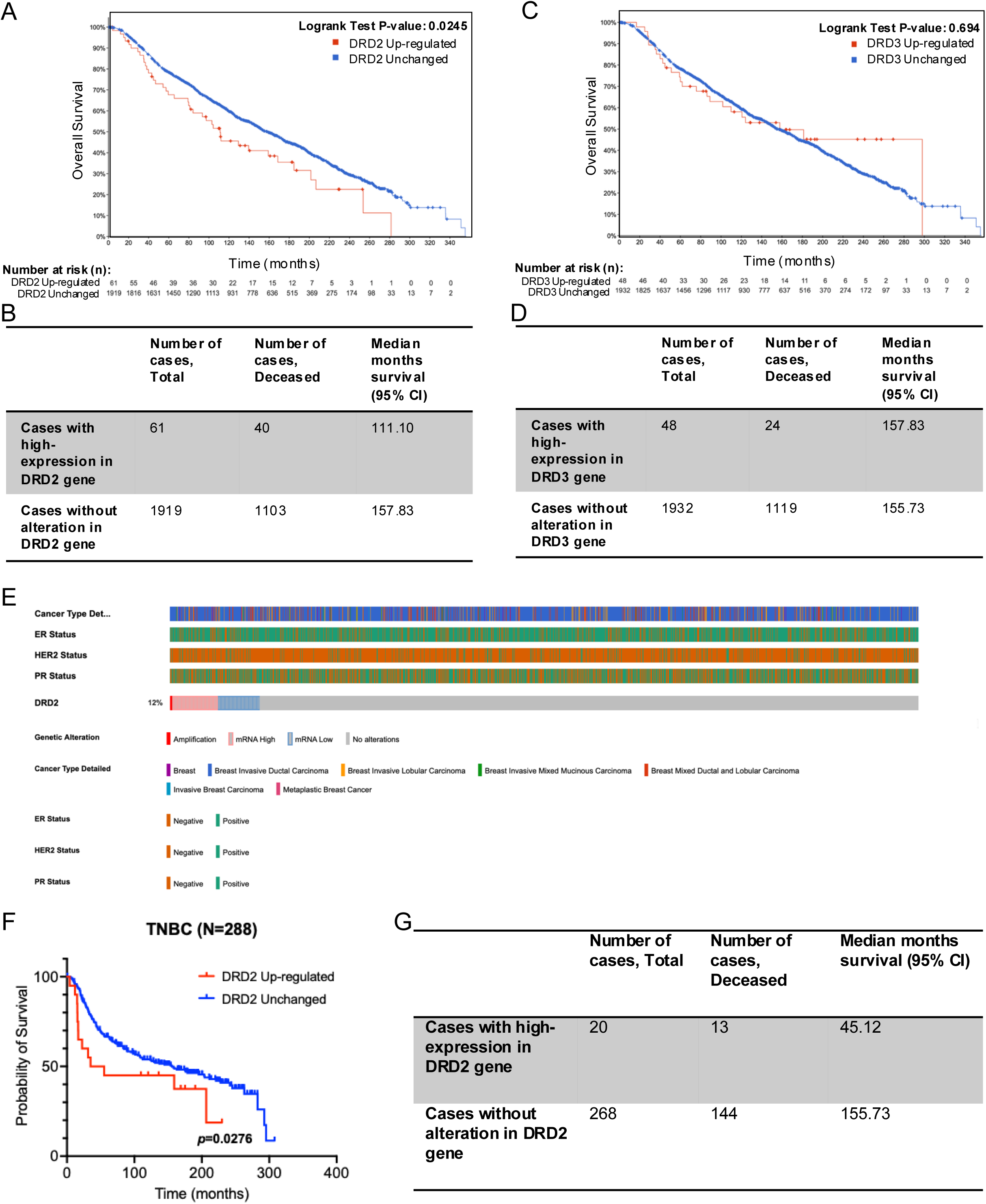
Evaluation of overall survival based on DRD2 and DRD3 signatures in invasive breast cancer patients. **(A)** Kaplan-Meier analysis of overall survival (OS) in invasive breast cancer patients stratified by DRD2 mRNA expression levels. The red line represents patients with upregulated DRD2 expression, while the blue line indicates those with unchanged expression. Patients in the DRD2 upregulated group exhibited significantly worse OS compared to the unchanged group (log-rank test, *p*=0.0245). The number of patients at risk is displayed below the Kaplan-Meier curve. **(B)** Summary table showing the total number of cases, deceased cases, and median survival of patients with either upregulated or unchanged DRD2 expression. **(C)** Kaplan-Meier analysis of OS in invasive breast cancer patients based on DRD3 mRNA expression levels. The red line denotes patients with upregulated DRD3 expression, while the blue line represents those with unchanged expression. OS in the DRD3 upregulated subgroup was not significantly different from the unchanged group (log-rank test, *p*=0.0694). The number of patients at risk is displayed below the Kaplan-Meier curve. **(D)** Summary table presenting the total number of cases, deceased cases, and median survival of patients with upregulated or unchanged DRD3 expression. **(E)** Heatmap summarizing breast-tumor cohorts: the top four tracks display histologic subtype and ER, HER2, and PR status for each sample, while the bottom track shows DRD2 alterations, revealing that 12 % of tumors harbor amplification or upregulated DRD2 expression. **(F)** Kaplan-Meier analysis of OS in TNBC patients stratified by DRD2 mRNA expression levels. The red line represents patients with up-regulated DRD2 expression, while the blue line indicates those with unchanged expression. OS in the DRD2 up-regulated subgroup was significantly different from the unchanged group (log-rank test, *p*=0.0276). **(G)** Summary table showing the total number of cases, deceased cases, and median survival of patients with either upregulated or unchanged DRD2 expression.

Given the heterogeneity of breast cancer and the aggressive nature of TNBC, we conducted a focused analysis to examine the prognostic value of DRD2 specifically within the TNBC subset. We filtered the TCGA dataset to include only patients with confirmed ER-negative, PR-negative, and HER2-negative status (N = 288). Patients were then stratified based on DRD2 expression using a more stringent Z-score cutoff of ±1.5 to better capture high-expression ones **(Figure 1E)**. Within this TNBC cohort, patients with upregulated DRD2 expression had a markedly poorer median OS of 45.12 months, compared to 155.73 months for patients with unchanged expression levels (*p* = 0.0276) **(Figure 1F/G)**. These findings indicate that elevated DRD2 expression may serve as a negative prognostic biomarker in TNBC, potentially identifying a high-risk subgroup of patients with worse clinical outcomes.

### Dopamine receptor expression in human triple-negative breast cancer cell lines under baseline and post-radiation conditions

We next examined the expression of dopamine receptors (DRD2 and DRD3) in established human TNBC cell lines. Cells were cultured both as monolayers and as mammospheres, the latter of which enriches for stem/progenitor cell populations. Interestingly, expression levels of DRD2 and DRD3 were significantly elevated in mammosphere cultures compared to their corresponding monolayer controls across all three TNBC lines tested (SUM159-PT, BT-549, and MDA-MB-231) (**Figure 2A**; black dashed line indicates baseline). These findings are consistent with previous reports showing that dopamine receptors, particularly DRD2 and DRD3, are broadly upregulated across multiple breast cancer subtypes, including lumina A, lumina B, HER2+-enriched non-luminal, and TNBC subtypes, relative to normal breast tissue [24].

**Figure 2.**
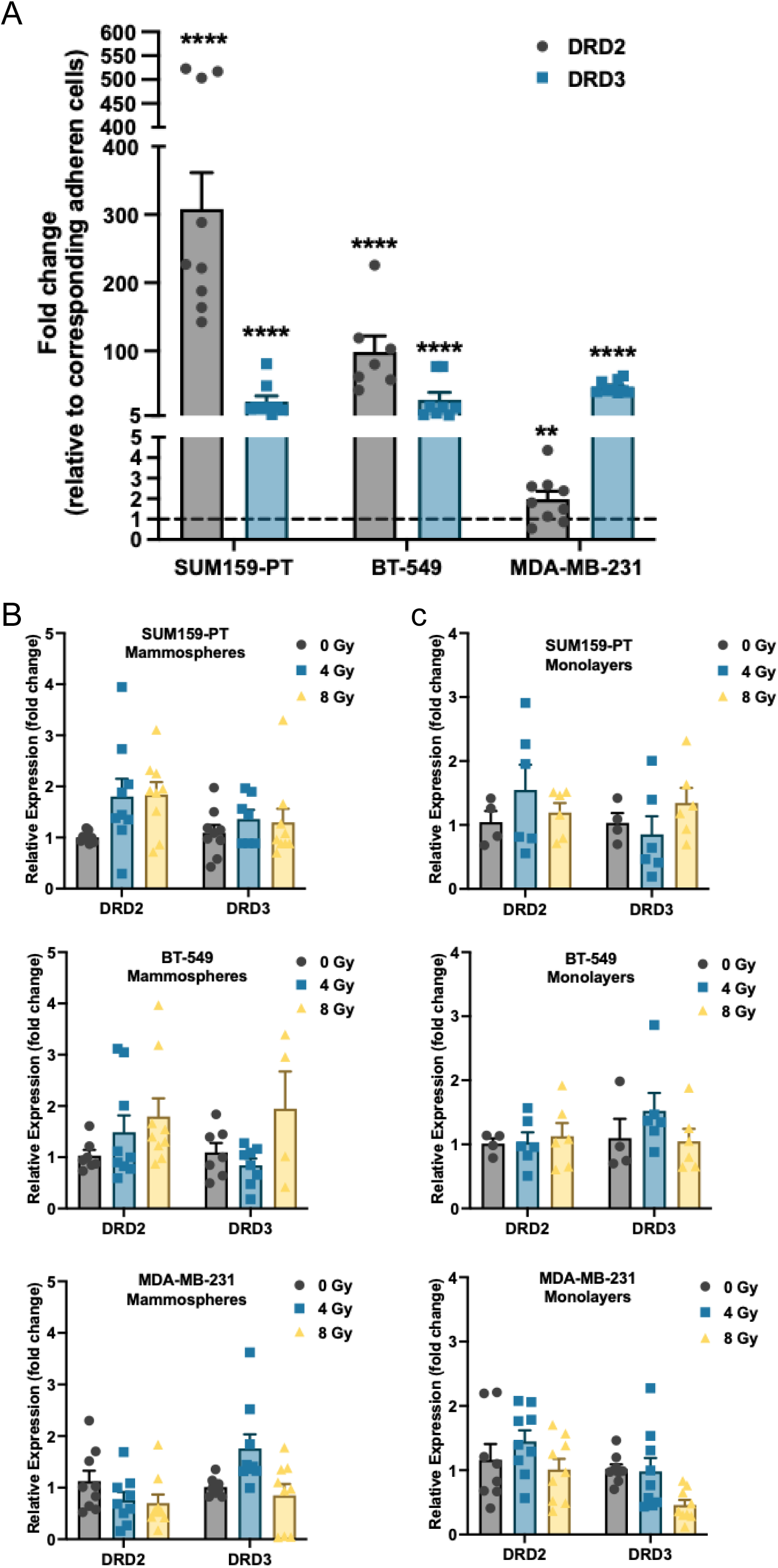
Dopamine receptor expression in human triple-negative breast cancer cell lines under baseline and post-radiation conditions. **(A)** Relative expression levels of DRD2 and DRD3 in mammospheres compared to their corresponding monolayers in SUM159-PT, BT-549, and MDA-MB-231 triple-negative breast cancer cell lines, as assessed by qRT-PCR. **(B/C)** DRD2 and DRD3 expression levels in mammospheres and monolayers of SUM159-PT, BT-549, and MDA-MB-231 following exposure to a single dose of 4 Gy or 8 Gy radiation, measured at 24 hours post-treatment. All experiments have been performed with at least 3 biological independent repeats. *P*-values were calculated using one-way ANOVA. ** *p*-value < 0.01, **** *p*-value < 0.0001.

Given that radiation can alter the surface expression of certain receptors, we also investigated DRD2 and DRD3 levels following exposure to single doses of 0, 4, or 8 Gy radiation under both adherent and mammosphere conditions. At 24 hours post-irradiation, DRD2 and DRD3 expression showed slight increases. However, these changes were not statistically significant **(Figure 2B)**. Together, these results suggest that DRD2 and DRD3 are preferentially expressed in TNBC stem/progenitor-like cell populations cultured under mammosphere conditions, and that their expression is relatively stable following sub-lethal radiation exposure. This stability may indicate a persistent receptor presence within radioresistant populations, highlighting a potential role for dopamine receptors in the maintenance of therapy-resistant TNBC cells.

### Quetiapine decreases clonogenicity and self-renewal in triple-negative breast cancer

To evaluate the potential anti-tumor effects of QTP in TNBC, we first performed clonogenic survival assays using monolayer cultures of all three TNBC lines (SUM159-PT, BT-549, and MDA-MB-231). Treatment with QTP resulted in a significant, dose-dependent reduction in colony formation capacity across all tested cell lines, and the effects seemed to reach a plateau at approximately 10 µM concentration. These findings strongly suggest that QTP has a robust inhibitory effect on the clonogenicity of tumor cells **(Figure 3A-D)**. Next, we examined whether QTP enhances cellular radiosensitivity by performing mammosphere-forming assays with increasing doses of radiation (0, 2, 4, 6, and 8 Gy) using SUM159-PT and BT-549 cells. Intriguingly, QTP treatment did not exhibit significant radiosensitizing effects, as the surviving fraction of mammosphere remained largely unaffected by combined radiation and QTP treatments compared to radiation alone **(Figure 3E/F)**. Microscopic examination of mammospheres revealed that QTP treatment resulted in substantially fewer and smaller mammospheres compared to DMSO or radiation alone groups **(Figure 3G)**. Additionally, extreme limiting dilution analyses demonstrated that QTP significantly reduced mammosphere-forming capacity and stem cell frequency, both in sham-irradiated cells and in cells receiving single doses of 4 or 8 Gy radiation **(Figure 3H-K)**. Taken together, our results demonstrate that while QTP strongly suppresses clonogenic growth and the self-renewal capacities of TNBC cells, it does not directly increase tumor cell radiosensitivity.

**Figure 3.**
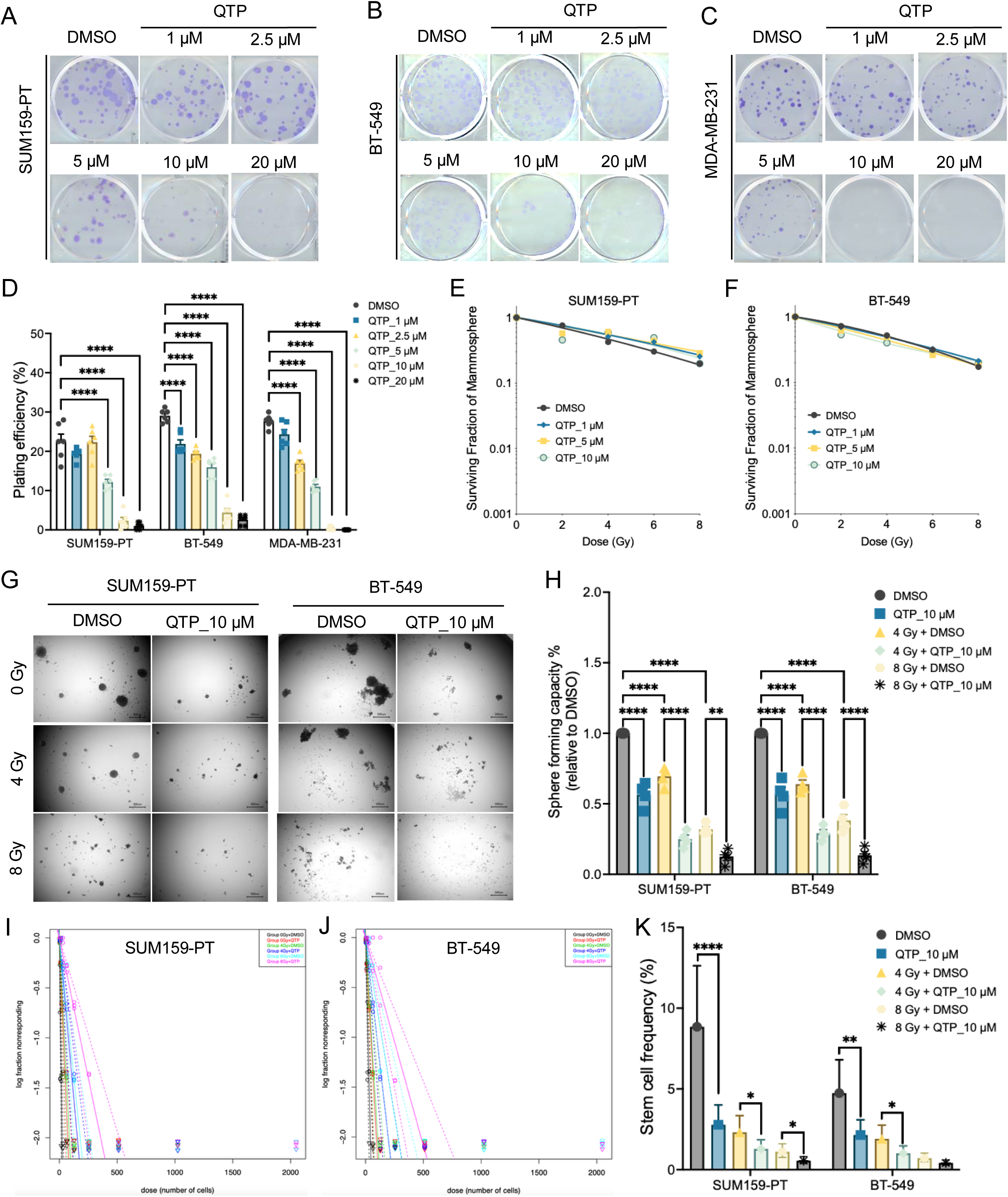
Quetiapine decreases clonogenicity and self-renewal in triple-negative breast cancer. **(A-C)** Clonogenic assays performed on monolayer cultures of patient-derived triple-negative breast cancer cells (SUM159-PT, BT-549, and MDA-MB231) treated with quetiapine at 1, 2.5, 5, 10, or 20 µM, or with DMSO (solvent control). **(D)** The colony number was counted and presented as the percentage relative to the initial number of cells plated. **(E/F)** Surviving fraction of SUM159-PT and BT-549 mammospheres treated with quetiapine at 1, 5, or 10 µM. **(G)** Representative images of SUM159-PT and BT-549 mammospheres treated with quetiapine (10 µM), with or without a single 4 Gy or 8 Gy radiation dose. **(H)** Mammosphere forming capacity of SUM159-PT and BT-549 spheres treated with quetiapine (10 µM), with or without a single 4 Gy or 8 Gy radiation dose. **(I/J)** Extreme Limiting Dilution Analysis (ELDA) plots showing the relationship between the number of cells plated per well and the fraction of negative wells (no colonies) in SUM159-PT and BT-549 mammospheres following the combination of radiation and quetiapine. **(K)** Estimated stem cell frequencies and 95% confidence intervals in each treatment group. All experiments have been performed with at least 3 biological independent repeats. *P*-values were calculated using one-way ANOVA. * *p*-value < 0.05, ** *p*-value < 0.01, *** *p*-value < 0.001, **** *p*-value < 0.0001.

### Quetiapine induces apoptosis and inhibits migratory capacity in TNBC cells

To elucidate the cellular mechanisms underlying the anti-tumor effects of QTP in TNBC cells, we cultured SUM159-PT and BT-549 cells in monolayers and treated them with 10 µM QTP, either alone or in combination with a single dose of 4 Gy radiation. Cell viability and proliferation were monitored at a regular interval (every 5 days) over a 20-day period. Notably, QTP treatment resulted in a significant reduction in cell proliferation at day 10 post-treatment in SUM159-PT cells **(Figure 4A)**, while in BT-549 cells, the inhibitory effects were pronounced as early as day 5 **(Figure 4B)**. Furthermore, combined treatment with QTP and radiation markedly reduced cell viability compared to radiation alone in both lines **(Figure 4CD)**.

**Figure 4.**
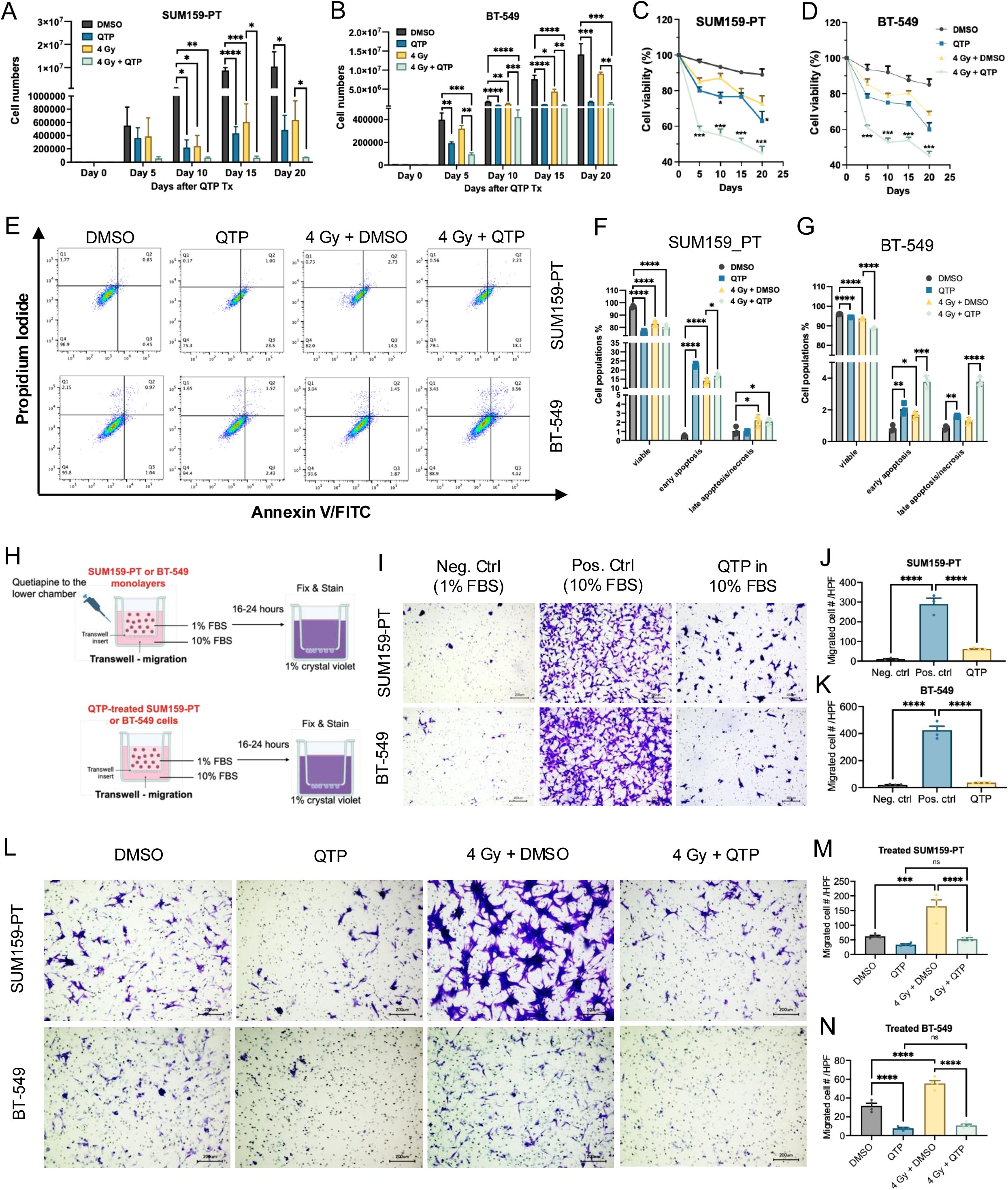
Quetiapine promotes apoptosis and reduces the migratory capacity of TNBC cells. **(A-D)** Quantification of cell number and viability in SUM159-PT and BT-549 cells treated with 10 µM QTP, with or without a single dose of 4 Gy radiation, assessed on days 5, 10, 15, and 20 post-treatment. **(E)** Representative Annexin V/PI dot plots of SUM159-PT and BT-549 cells treated with QTP (10 µM), with or without a single 4 Gy dose of radiation, analyzed 72 hours post-treatment. **(F/G)** Quantification of viable, early apoptotic, and late apoptotic/necrotic cell populations. **(H)** Schematic illustration of two transwell migration assay setups. **(I-K)** Transwell invasion assay of SUM159-PT and BT-549 cells with QTP (10 µM) added to the lower chamber of an 8-µm pore insert. Media containing 1% FBS and 10% FBS served as negative and positive control, respectively. Migrated cells were quantified per high-power field using Image J. **(L-N)** Invasion assay of SUM159-PT and BT-549 cells pretreated with QTP (10 µM) for 48 hours, with or without a single dose of 4 Gy radiation, followed by migration analysis. All experiments have been performed with at least 3 biological independent repeats. *P*-values were calculated using one-way ANOVA. * *p*-value < 0.05, ** *p*-value < 0.01, *** *p*-value < 0.001, **** *p*-value < 0.0001, ns: not significant.

To determine whether the observed inhibition of cell proliferation resulted from induction of cell apoptosis, we conducted Annexin V/PI staining followed by flow cytometry analysis. Our results revealed that QTP treatment significantly reduced the percentage of viable cells and markedly increased the proportion of early apoptotic cells (defined as the Annexin V-positive/PI-negative). Additionally, there was a substantial increase in late apoptotic or necrotic populations (Annexin V-positive/PI-positive) **(Figure 4E-G)**.

TNBC is known for its aggressive behavior, often spreading more rapidly and frequently than other breast cancer subtypes, which leads to a high risk of early metastasis. To investigate whether QTP impacts the migratory/invasive properties of TNBC cells, we performed transwell migration assay using two experimental approaches **(Figure 4H)**. In the first setup, QTP was added to the lower chamber to assess its effect as a chemoattractant inhibitor. Here, we demonstrated that QTP significantly reduced the migratory capacity of both SUM159-PT and BT-549 cells, as evidenced by a decreased number of cells penetrating through the transwell membrane **(Figure 4I-K)**. In the second approach, cells were pre-treated with QTP alone or in combination with a single dose of 4 Gy radiation, prior to seeding. The pre-treated cells displayed markedly impaired migration capacity compared to untreated and radiation-only controls **(Figure 4L-N)**. Although the difference between QTP alone and QTP combined with radiation did not reach statistical significance, our primary objective was to highlight the increased motility observed in cells that survive irradiation, a phenomenon consistent with prior studies reporting radiation-induced enhancement of migratory and invasive behavior in cancer cells [25, 26]. Notably, the addition of QTP appeared to attenuate this radiation-associated increase in motility, suggesting a potential inhibitory effect of QTP on radiation-enhanced cell movement. Overall, these observations may have biological relevance in the context of tumor recurrence and metastasis.

### Quetiapine induces DNA double-strand breaks and impairs DNA repair in TNBC cells

Based on our observations that QTP reduces cell viability, impairs self-renewal, and suppresses migratory potential in TNBC cells, we next sought to explore whether its anti-tumor effects are associated with the induction of genomic instability and disruption of DNA repair mechanisms. Given that radiation therapy primarily exerts its cytotoxic effects through the induction of DNA double-strand breaks (DSBs), and the ability to repair such damage is essential for tumor cell survival and therapy resistance. In order to examine whether QTP contributes to DNA damage or interferes with the repair process, we evaluated the formation and resolution of γ-H2AX foci, a widely accepted marker of DSBs, at 6 hours and 24 hours following exposure to a single dose of 4 Gy radiation, in the presence or absence of QTP using SUM159-PT cells. At 6 hours post-radiation, QTP-treated cells exhibited a marked increase in γ-H2AX positive cells compared to both DMSO control and cells treated with radiation alone, suggesting that QTP enhances the induction of DNA damage in TNBC cells **(Figure 5A/B)**. Notably, while radiation-induced γ-H2AX foci typically resolve by 24 hours post-radiation as a result of efficient DNA repair, QTP-treated cells displayed a persistent γ-H2AX signal at this later time point **(Figure 5C/D)**. This indicates a significant delay in DNA repair and suggests that QTP may impair the resolution of DSBs.

**Figure 5.**
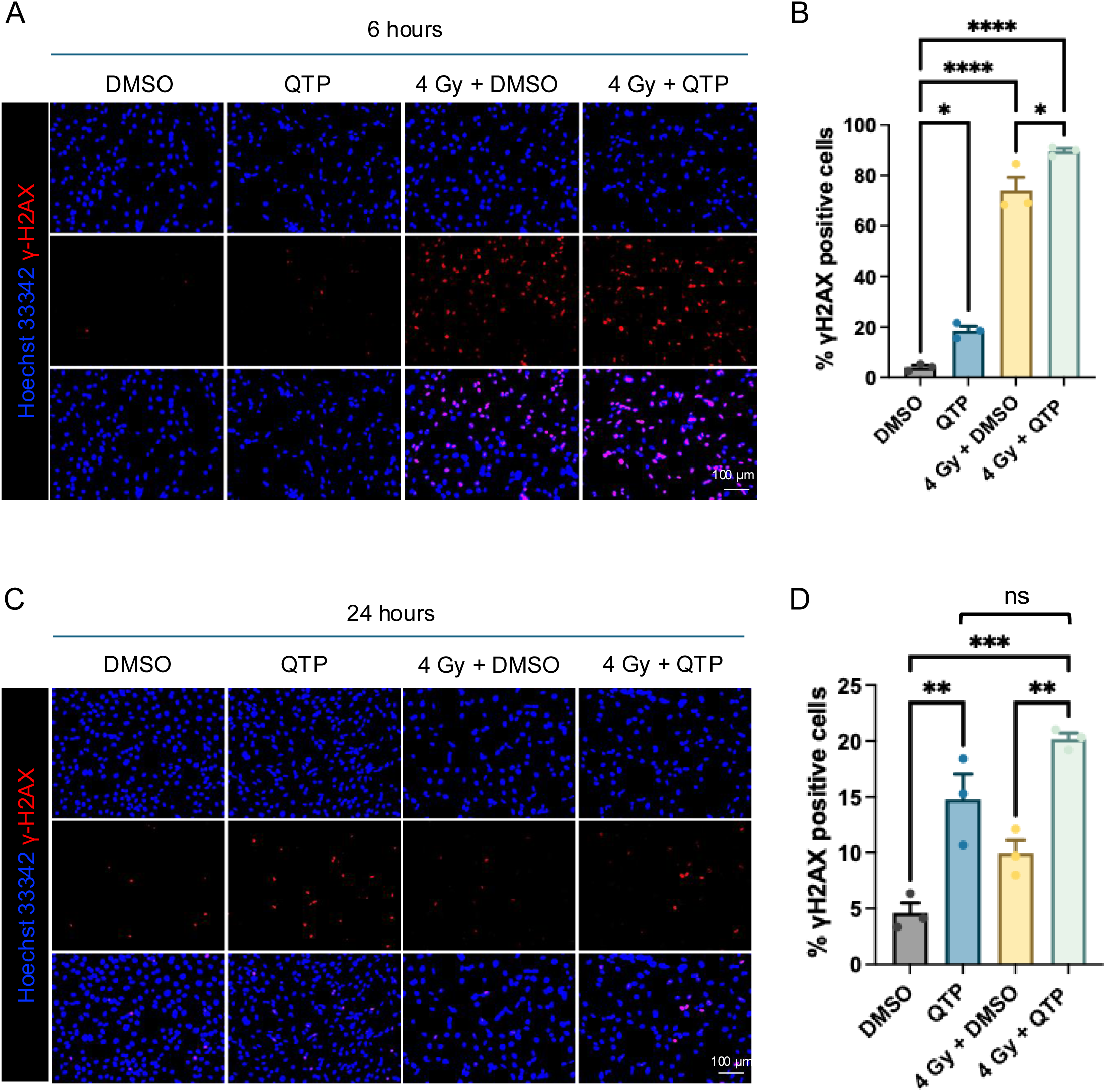
Quetiapine induces DNA double-strand breaks and delays DNA repair in TNBC cells. **(A)** Representative immunofluorescence images of γ-H2AX staining in SUM159-PT cells treated with 10 µM QTP, with or without a single dose of 4 Gy radiation, analyzed 6 hours post-radiation. **(B)** Quantification of γ-H2AX-positive cells per high-power field using Image J. **(C/D)** Representative γ-H2AX immunofluorescence images and corresponding quantification at 24 hours post-radiation under the same treatment conditions. All experiments have been performed with at least 3 biological independent repeats. *P*-values were calculated using one-way ANOVA. * *p*-value < 0.05, ** *p*-value < 0.01, *** *p*-value < 0.001, **** *p*-value < 0.0001, ns: not significant.

### Anti-tumor effects of QTP and chemotherapeutic agents in TNBC cells

To evaluate the potential of QTP as a combination therapy in TNBC, we investigated whether it could enhance the efficacy of standard chemotherapeutic agents commonly used in clinical settings. Specifically, we tested QTP in combination with doxorubicin (DOX), paclitaxel (TAXOL), and 5-fluorouracil (5-FU) in SUM159-PT cells. Given the high potency of QTP at 10 µM, we reduced its concentration to 5 µM to better assess synergistic interactions. Clonogenic survival assays revealed a substantial reduction in colony-forming ability when QTP was combined with low doses of 5-FU (0.5 µM) **(Figure 6A)** or TAXOL (0.5 µM) **(Figure 6B)**, indicating enhanced cytotoxicity beyond single-agent treatments. Notably, the combination of QTP and DOX demonstrated a clear dose-dependent suppression of clonogenicity **(Figure 6C)**.

**Figure 6.**
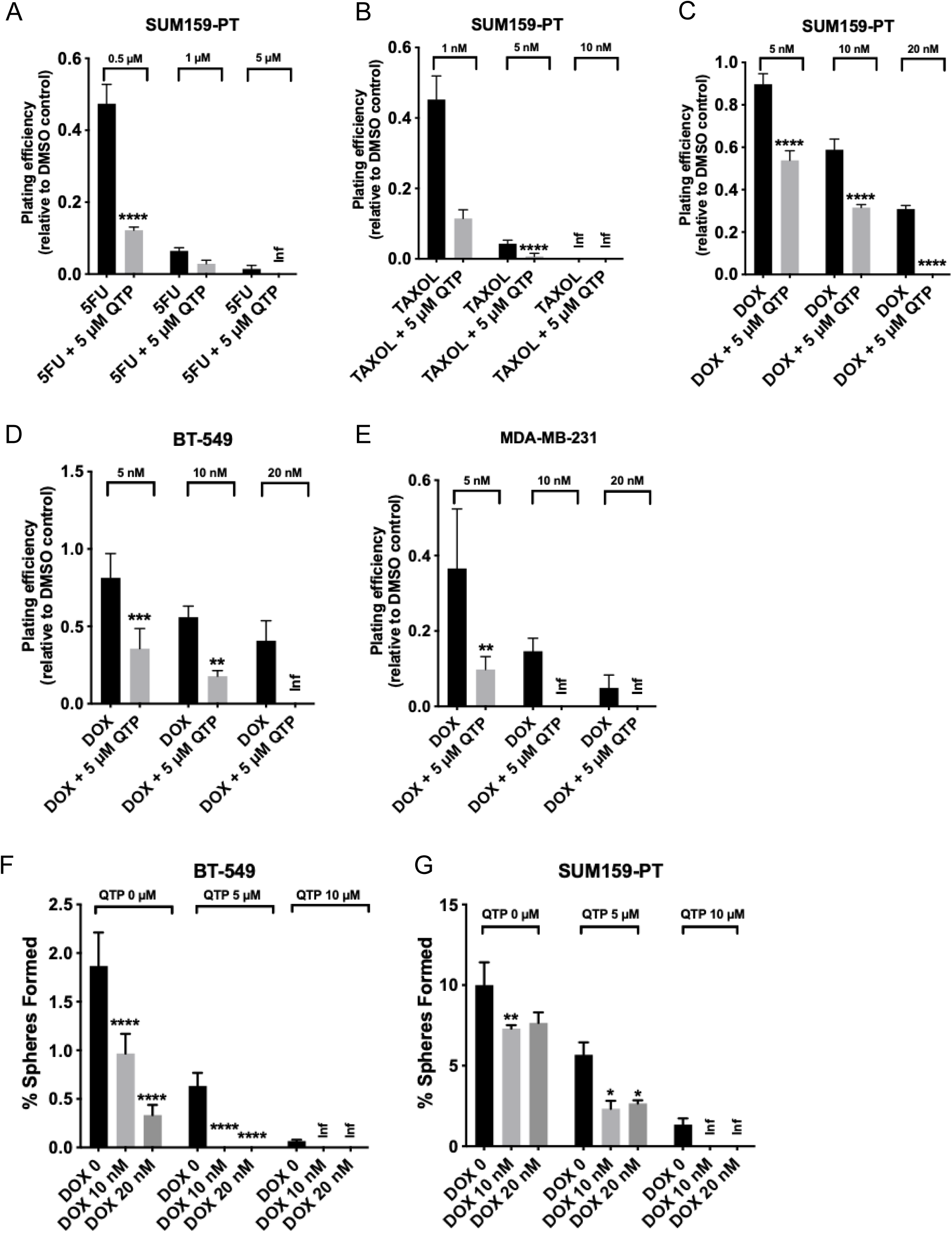
Anti-tumor effects of quetiapine and chemotherapeutic agents in TNBC cells. **(A-C)** Quantification of plating efficiency in SUM159-PT cells treated with 5 µM QTP in combination with increasing concentrations of 5-fluorouracil (5-FU), paclitaxel (TAXOL), or doxorubicin (DOX), assessed via clonogenic survival assays. **(D/E)** Clonogenic survival of BT-549 and MDA-MB-231 cells treated with 5 µM QTP in combination with 5 nM, 10 nM, or 20 nM DOX. **(F/G)** Mammosphere-forming assays of BT-549 and SUM159-PT cells treated with varying doses of QTP in combination with DOX to evaluate effects on self-renewal. All experiments have been performed with at least 3 biological independent repeats. *P*-values were calculated using two-way ANOVA. * *p*-value < 0.05, ** *p*-value < 0.01, *** *p*-value < 0.001, **** *p*-value < 0.0001, inf: infinity.

To confirm the broader applicability of this synergy, we extended the analysis to two additional TNBC cell lines, BT-549 and MDA-MB-231. In BT-549 cells, the combined treatment of 5 µM QTP with 20 nM DOX eliminated colony formation **(Figure 6D)**. Similarly, MDA-MB-231 cells showed a near-complete loss of clonogenic potential when treated with 5 µM QTP in combination with either 10 nM or 20 nM DOX **(Figure 6E)**. These consistent findings across multiple TNBC lines support the conclusion that QTP significantly sensitizes TNBC cells to doxorubicin and other chemotherapeutic agents.

Furthermore, we examined whether this synergistic interaction extended to the self-renewal capacity of TNBC cells using the mammosphere-forming assay. The combination of QTP and DOX completely abolished sphere formation in BT-549 cells **(Figure 6F)**, and similar effects were observed in SUM159-PT cells treated with 10 µM QTP plus DOX **(Figure 6G)**. These results indicate that QTP not only enhances the cytotoxic effects of chemotherapy on bulk tumor cells but also effectively targets cancer stem-like cells, supporting its potential as a valuable adjuvant to conventional chemotherapy in the treatment of TNBC.

## Discussion

The psychiatric drug quetiapine, a second-generation atypical antipsychotic commonly used for the treatment of schizophrenia and bipolar disorder, has recently demonstrated promising anti-tumor effects in several preclinical cancer models [21, 27]. Despite these encouraging results, its therapeutic efficacy has not yet been evaluated in triple-negative breast cancer, a highly aggressive subtype characterized by limited treatment options and poor patient prognosis. In this study, we report, for the first time, that QTP not only inhibits tumor clonogenicity and self-renewal but also enhances the efficacy of both radiation and chemotherapeutic agents in multiple TNBC cell models.

More importantly, these effects are observed when QTP is used alone and are further amplified in combination with clinically relevant treatments, such as ionizing radiation, doxorubicin, paclitaxel, and 5-fluorouracil. This suggests a synergistic or additive interaction between QTP and standard-of-care therapies, potentially offering a new combination strategy for this highly aggressive subtype. Mechanistically, we show that QTP induces apoptosis and promotes the accumulation of DNA double-strand breaks. Notably, QTP also delays the resolution of γ-H2AX foci over time, suggesting that it disrupts the DNA damage response and impairs DNA repair process. This is particularly significant in the context of radiotherapy or genotoxic chemotherapy, where DNA repair pathways are critical for tumor cell survival [28, 29]. Additionally, QTP suppresses the bulk tumor cell migration, as well as the cells that survive sublethal radiation, indicating its potential to prevent metastatic progression and recurrence.

These observations are in line with previous reports on the anti-tumor activity of dopamine receptor antagonists [15, 30, 31]. Prior studies have demonstrated that other antipsychotic agents such as thioridazine and domperidone could target cancer stem-like cells and reduce self-renewal in TNBC via DRD2 inhibition and subsequent suppression of downstream pathways such as STAT3 and JAK/STAT signaling [30, 31]. Our findings extend these insights by identifying QTP, a DRD2/DRD3 antagonist with a well-established clinical safety profile, as a novel agent capable of replicating these effects while offering additional therapeutic advantages. Unlike previous studies that focused solely on monotherapy, our work highlights the potential of QTP as a promising adjuvant therapeutic agent when combined with radiation and chemotherapy.

While some of our assays did not show statistically significant differences between QTP alone and QTP combined with 4 Gy after normalizing to the DMSO control, we believe the findings are still biologically meaningful. These experiments were designed to explore how cells behave after surviving sub-lethal radiation. As reported in previous studies, we observed that radiation alone tends to increase TNBC cell motility, reflecting a known pro-migratory effect of radiation. Interestingly, this increase was reduced when cells were treated with QTP, suggesting that QTP may help suppress radiation-induced cell movement. The absence of an additive effect with the combination treatment could mean that QTP alone is already achieving near-maximal inhibition, or that QTP and radiation affect similar pathways. Overall, these observations highlight the value of considering biological trends even when statistical differences are modest, especially when studying therapy resistance. Additional mechanistic work and extended time-point analyses will be important to better understand how QTP interacts with radiation over time.

TNBC patients have a higher likelihood of developing brain metastases compared to those with hormone receptor-positive subtypes. These metastases not only occur more frequently but also tend to develop earlier in the course of the disease [32]. Once brain involvement is detected, the prognosis is generally poor, with median overall survival typically ranging from 4 to 9 months, making it a major driver of morbidity and mortality in patients with metastatic TNBC [33–35]. Beyond the physical challenges posed by metastasis, many TNBC patients also suffer from the neurological and psychological side effects of systemic chemotherapy, including cognitive dysfunction (“chemo brain”), anxiety, depression, and sleep disturbances. These symptoms can severely compromise quality of life and hinder adherence to treatment regimens [36–38]. Here QTP presents as a compelling candidate for repurposing in oncology due to several favorable pharmacological properties. It readily crosses the blood-brain barrier, has a well-documented safety profile with a low incidence of severe side effects, and is widely used in clinical practice for managing psychiatric conditions. In previous studies, QTP has been shown to effectively eliminate glioma-initiating cells, highlighting its capacity to act against therapy-resistant cancer stem-like cells within the brain [21]. These findings raise the possibility that QTP may also exhibit therapeutic efficacy against TNBC brain metastases. This multifunctional profile could be advantageous in managing the complex needs of TNBC patients with central nervous system (CNS) involvement, and it warrants further investigation in preclinical and clinical settings.

Despite these promising findings, our study has several limitations. Most notably, all experiments were conducted *in vitro*, and the complexities of the tumor microenvironment, drug metabolism, and immune interactions could potentially influence its therapeutic efficacy *in vivo*. Future studies are needed to evaluate its efficacy in TNBC xenograft or patient-derived orthotopic mouse models to confirm its anti-tumor and anti-metastatic potential in a physiologically relevant context. Furthermore, although QTP is considered to act as a DRD2/DRD3 antagonist, it interacts with several other receptors (e.g., serotonin and adrenergic receptors), and their contributions to the observed phenotypes remain to be explored. Lastly, the molecular pathways downstream of DRD2/DRD3 that mediate QTP’s effects in TNBC warrant further mechanistic study.

## Conclusion

Our study provides new insight into the anti-tumor properties of QTP in TNBC and identifies it as a promising candidate for combination therapies. By impairing DNA repair, inducing apoptosis, and suppressing migratory and stem-like features of TNBC cells, QTP has the great potential to overcome some of the major therapeutic challenges in this aggressive breast cancer subtype. These findings lay the groundwork for future preclinical and clinal studies aimed at repurposing QTP as an adjuvant treatment strategy for TNBC, potentially improving patient outcomes through a more cost-effective, well-tolerated therapeutic approach.

## List of Abbreviations

TNBC: triple-negative breast cancer
QTP: quetiapine
ER: estrogen receptor
PR: progesterone receptor
HER2: human epidermal growth factor receptor 2
DRD2: dopamine receptors D2
DRD3: dopamine receptors D3
DRD4: dopamine receptors D4
PI: propidium iodide
TAXOL: paclitaxel
DOX: doxorubicin
5-FU: 5-fluorouracil
DSBs: DNA double-strand breaks
CNS: central nervous system
OS: overall survival
ELDA: Extreme Limiting Dilution Analysis
TCGA: The Cancer Genome Atlas.

## Data Availability

Research data are stored in an institutional repository and will be shared upon request to the corresponding author.

## Acknowledgements

Not applicable.

## Supplementary Materials

Supplemental Figure 1: Evaluation of overall survival based on DRD2 and DRD3 signatures in invasive breast cancer patients.

Supplemental Table 1: PCR primer sequences (Integrated DNA Technologies) used for RT-PCR experiments.

