## Supplemental Figure 1 for "Repurposing Quetiapine as an Adjuvant Therapeutic Agent for Triple-Negative Breast Cancer"

### Slide 1
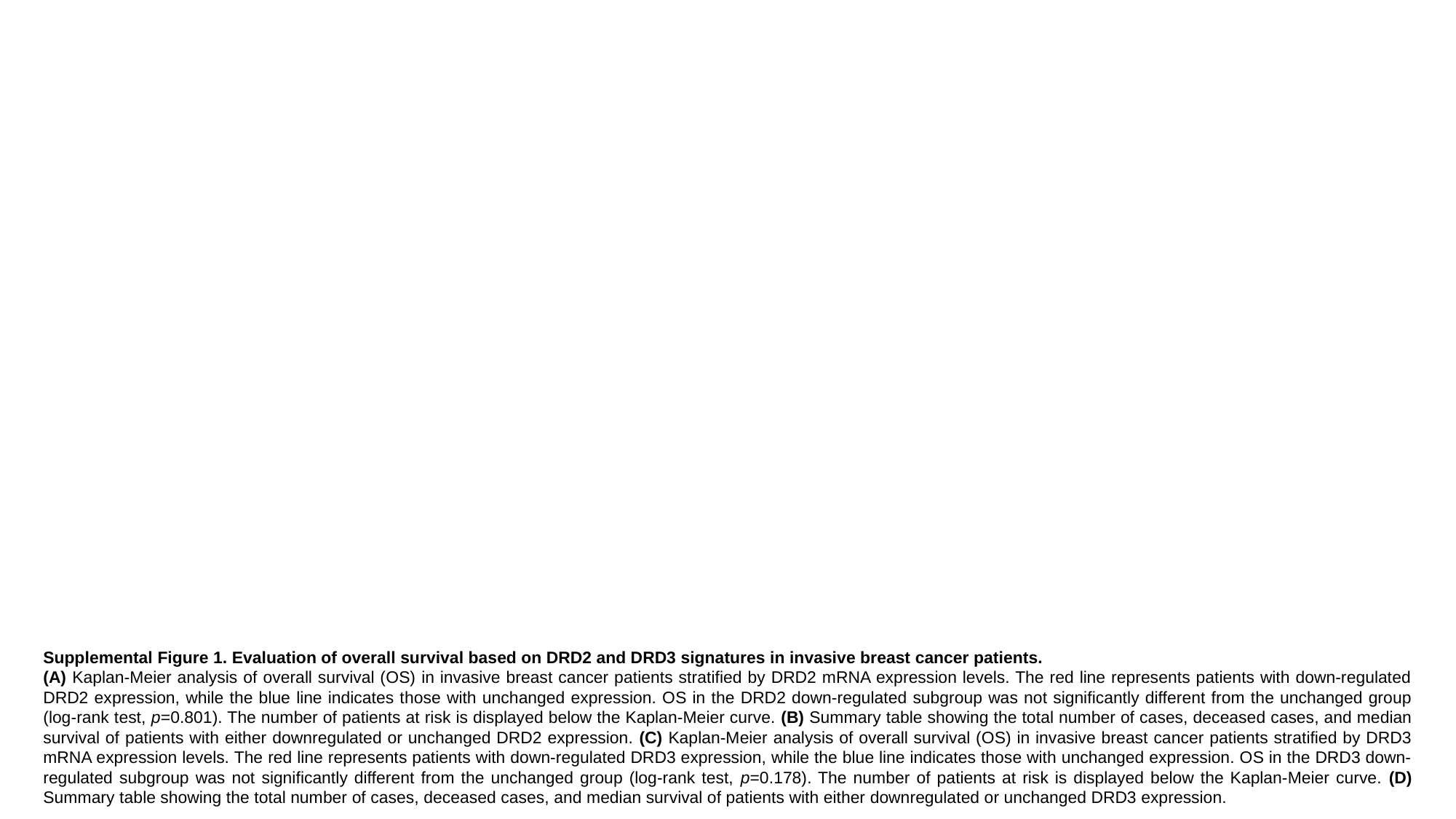

Supplemental Figure 1. Evaluation of overall survival based on DRD2 and DRD3 signatures in invasive breast cancer patients.
(A) Kaplan-Meier analysis of overall survival (OS) in invasive breast cancer patients stratified by DRD2 mRNA expression levels. The red line represents patients with down-regulated DRD2 expression, while the blue line indicates those with unchanged expression. OS in the DRD2 down-regulated subgroup was not significantly different from the unchanged group (log-rank test, p=0.801). The number of patients at risk is displayed below the Kaplan-Meier curve. (B) Summary table showing the total number of cases, deceased cases, and median survival of patients with either downregulated or unchanged DRD2 expression. (C) Kaplan-Meier analysis of overall survival (OS) in invasive breast cancer patients stratified by DRD3 mRNA expression levels. The red line represents patients with down-regulated DRD3 expression, while the blue line indicates those with unchanged expression. OS in the DRD3 down-regulated subgroup was not significantly different from the unchanged group (log-rank test, p=0.178). The number of patients at risk is displayed below the Kaplan-Meier curve. (D) Summary table showing the total number of cases, deceased cases, and median survival of patients with either downregulated or unchanged DRD3 expression.
