## Supplemental Table 1 for "Repurposing Quetiapine as an Adjuvant Therapeutic Agent for Triple-Negative Breast Cancer"

**Supplemental Table 1. PCR primer sequences (Integrated DNA Technologies) used for RT-PCR experiments.**

| Species | Gene name | Primer sequence (5’–3’) |
| --- | --- | --- |
| Human | DRD2 | Forward: TGTACAATACGCGCTACAGCTCCA  Reverse: ATGCACTCGTTCTGGTCTGCGTTA |
| Human | DRD3 | Forward: GTGGTGTCCTTCTACCTGCC  Reverse: GAGAGAGGGTTTGTTGGGGG |
| Human | PPIA | Forward: ATGCTGGACCCAACACAAAT  Reverse: TCTTTCACTTTGCCAAACACC |
